# BrightEyes-FFS: an open-source platform for comprehensive analysis of fluorescence fluctuation spectroscopy experiments with small detector arrays

**DOI:** 10.64898/2026.04.08.717207

**Authors:** Eli Slenders, Eleonora Perego, Sabrina Zappone, Giuseppe Vicidomini

## Abstract

Fluorescence fluctuation spectroscopy (FFS) is an ensemble of techniques for quantitative measurement of molecular dynamics and interactions. Recently, the introduction of small-format array detectors has opened up a new range of spatiotemporal information, allowing for more detailed analysis of system kinetics. However, there is currently no open-source software available for analyzing the high-dimensional FFS data sets. We present BrightEyes-FFS, an open-source Python-based environment for FFS analysis with array detectors. The environment includes a Python package for reading raw FFS data, computing auto- and cross-correlations using various algorithms, and fitting the correlations to several models. A graphical user interface (GUI), available as a standalone executable, makes the analysis fast and user-friendly. An automated Jupyter Notebook writing tool enables transition from the GUI to Jupyter Notebook for custom analysis. We believe that BrightEyes-FFS will enable a wider community to study diffusion, flow, and interaction dynamics.

## 1. Introduction

Fluorescence fluctuation spectroscopy (FFS) is a family of powerful techniques for investigating molecular concentration, dynamics, interactions, confinement, and aggregation in complex environments such as living cells [1, 2, 3, 4]. In confocal-based fluorescence correlation spectroscopy (FCS), an excitation laser beam is focused by the objective lens and, combined with confocal detection, forms a diffraction-limited detection volume at a static position in the sample. Fluorescence from emitters inside the observation volume is measured with a point-detector. Spontaneous deviations from equilibrium, *e*.*g*., due to Brownian motion of the fluorophores, lead to temporal fluctuations in the fluorescence trace *I*(*t*). Calculating the autocorrelation function (ACF) *G*(*τ*) of the intensity trace over a range of lag times *τ* and fitting *G*(*τ*) to a model provides information on the sys-tem kinetics, such as the molecular diffusion coefficient, provided that the dimensions of the focal volume are known.

The confocality ensures a femtoliter-scale detection volume, which is required to restrict the number of molecules to only a few at the typical concentrations encountered in live-cell experiments, thereby enabling robust correlation analysis. In addition, the laser-scanning architecture of a confocal microscope allows (i) multiplexing the FCS experiment by acquiring data at different positions in the sample and (ii) scanning the focused laser on the sample during the acquisition, thus performing scanning FCS or image scanning FCS [5].

However, confocal-based FCS with a single-element detector has limitations in terms of detecting diffusion anisotropy or flow, such as superdiffusion caused by active transport or subdiffusion caused by molecular crowding or attractive interactions. The recent development of small detector arrays has significantly increased the amount of information that can be extracted from a single FFS measurement [6, 7, 8]. These detector arrays, such as fiber bundle-based photomultiplier arrays [9] and asynchronous readout single-photon avalanche diode (SPAD) arrays [10, 11], typically comprise a few tens of elements and enable imaging of the probed region with (sub-)microsecond temporal resolution, Fig. 1 (a, b). This newly accessible spatiotemporal information allows for detailed analysis of molecular mobility and diffusion modalities, including anomalous diffusion associated with biomolecular condensate formation [12]. Additionally, these detectors facilitate the investigation of flow dynamics, such as the direction and velocity of convective flow [13]. Apart from the increased information on the dynamics properties of the sample, the use of array detectors offers an additional technical advantage. Since the lateral displacement of each detector element relative to the central element is known, the separation between detection volumes is fixed a priori, eliminating the need for a separate calibration measurement of the observation volume [14].

**Figure 1:**
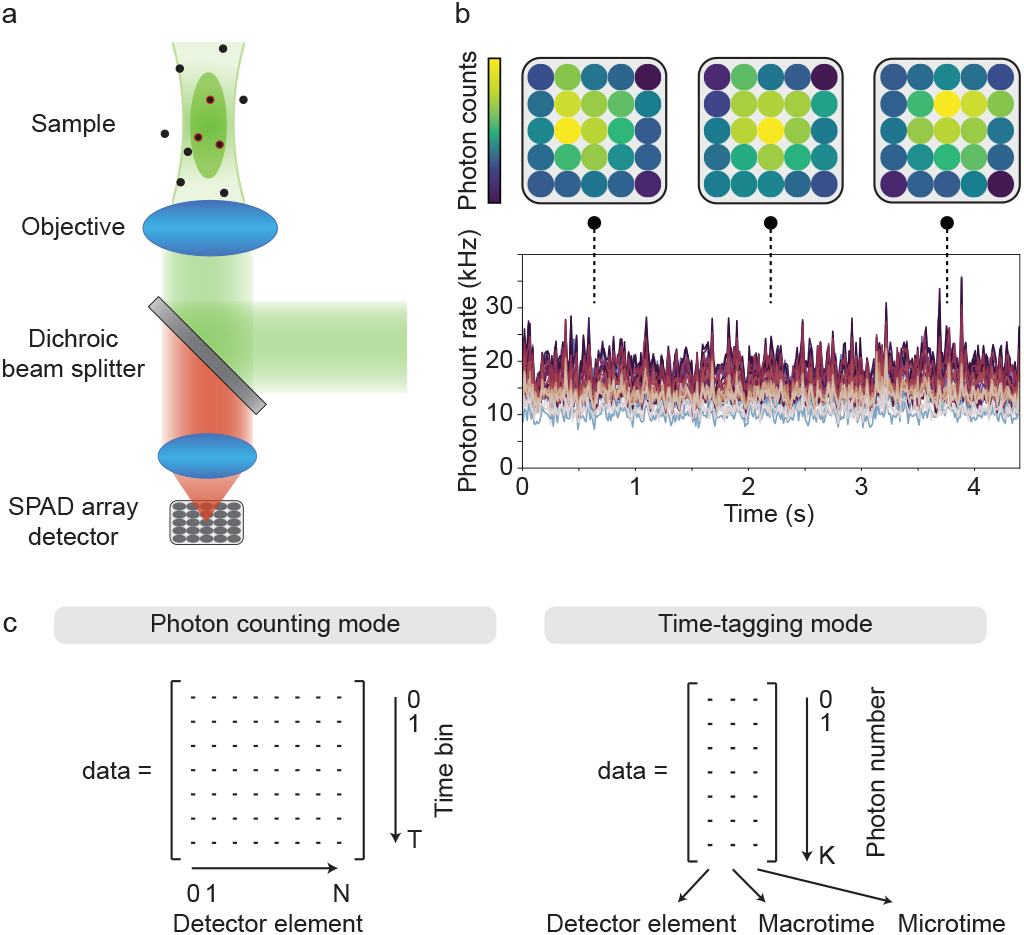
(a) Principle of FFS. A small array detector, here 5 × 5 detector elements, images the fluorescence fluctuations caused by fluorophores moving through the focal volume produced by an objective lens. (b) Illustrative FFS data set acquired with an array detector. Different colors in the time trace represent different detector elements. (c) Depending on the acquisition mode, a raw FFS data set can take different forms: (i) in photon counting mode, the data are a 2D array with for each time bin and detector element a given number of photons; (ii) in time-tagging mode, the data are a list with for each detector element the arrival macro- and microtimes for each photon.

Array detectors are becoming increasingly available and implemented in laser scanning microscopes (LSMs), *e*.*g*., the AiryScan from Zeiss, NSPARC from Nikon, MATRIX from Abberior, PDA-23 from PicoQuant’s Luminosa system, and PRISM from Genoa Instruments. The use of small array detectors for FFS necessitates the development of advanced data analysis protocols and software capable of calculating and fitting the numerous auto- and cross-correlation functions generated from a single FFS measurement. However, the lack of an open-source and user-friendly platform for such analysis currently restricts the accessibility of this technique to microscopists with good programming skills. To address these challenges, we developed BrightEyes-FFS, a Python-based open-source platform for array-detector based FFS analysis, including a graphical user interface (GUI) with an automated Jupyter Notebook writing tool.

## 2. Methods

Depending on the detector type and measurement protocol, an FFS experiment may yield a different data set. For analog array detectors or digital array detectors in photon counting mode, the data set comprises an intensity trace or photon count trace *I*_*c*_[*t*_*n*_], respectively, for each detector element *c* and discrete time bin *t*_*n*_ = *n*Δ*t*, Fig. 1 (c). For array detectors connected to a multichannel time-correlated single-photon counting (TCSPC) platform, the data is a list of photon arrival times *t*_*c*_(*i*), with *t* the (macro)time of the *i*-th photon in channel *c*, Fig. 1 (c). In the case of scanning-FFS, each time point is linked with an (x,y) position of the moving excitation beam. In this section, we show some examples of the variety of auto- and cross-correlations that can be calculated to analyze an FFS data set.

### 2.1 Conventional FCS

The most straightforward analysis is to treat the data from each detector element as an independent FCS measurement (*e.g*., for a 5 × 5 array detector, 25 autocorrelation curves can be calculated). For time-trace data, the correlation function for detector element *c* is calculated as:

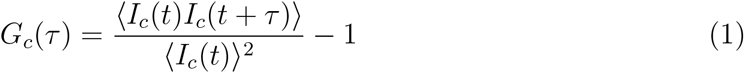

In practical implementations, such as the Python package Multipletau [15], correlations are computed on a logarithmic time scale by binning the data in increasingly larger time bins for increasing *τ*, thereby also speeding up the calculation of *G*(*τ*) for large *τ*. Alternatively, the autocorrelation can be calculated in Fourier space using the Wiener-Khinchin theorem, and selecting correlation values at logarithmically spaced *τ* values afterwards. The latter approach is particularly useful for orbital scanning-FCS, in which the excitation beam is moving in a small circular pattern, and the resulting oscillating behavior of the correlation curve would be smoothed out by the time binning with Multipletau. For TCSPC data, the correlation curve can be directly calculated on the raw data using the time-tag-to-correlation algorithm proposed by Wahl *et al*. [16].

Many models exist to fit the experimental correlation curve. In conventional FCS, and under the assumption of a 3D Gaussian focal volume with 1/*e*^2^ lateral waist *ω*_0_ and axial extension *z*_0_ and in the absence of triplet blinking, the ACF of a single freely diffusing component in 3*D* is:

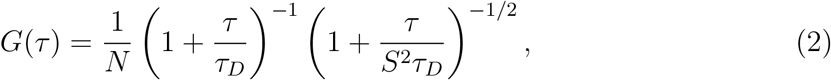

with 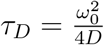 the diffusion time,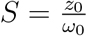 the shape parameter, *N* the apparent number of molecules in the focal volume, and *D* the diffusion coefficient.

With an array detector, the point-spread-function (PSF) for each element is the product of the excitation PSF *h*_1_ and the element specific detection PSF *h*_2,*d*_. All detection PSFs are identical, except for a lateral shift reflecting the position of the element in the array. Assuming *h*_1_ and *h*_2,*d*_ to be Gaussian functions with equal lateral waist, the product *h*_1_*h*_2,*d*_ is a Gaussian function with waist 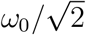, centered between *h*_1_ and *h*_2,*d*_. However, the Gaussian approximation is typically only valid for elements close to the optical axis [17]. Therefore, it is not recommended to use Eq. 2 for fitting ACFs calculated for elements near the edge of the detector.

Converting the fit parameters *N* and *τ*_*D*_ to the particle concentration and diffusion coefficient requires the confocal volume parameters *ω*_0_ and *S* to be known. The calibration of these parameters typically involves an FCS measurement of a sample of known diffusion parameters. Alternatively, in orbital scanning FCS, the excitation beam is moved rapidly in a small circular pattern and the beam waist can be derived from the measurement itself, provided that the orbit time is much shorter than the diffusion time [7].

By treating the data as a set of *N*_*c*_ autocorrelation curves, with *N*_*c*_ the number of detector elements, and fitting all curves independently from each other, all spatial information contained in the data set is discarded. Spot-variation FCS (svFCS) and cross-correlation FFS are analysis methods that exploit spatial information, providing information not only on the diffusion coefficient but also on the diffusion mode, flow parameters, and diffusion anisotropy.

### 2.2. Spot-variation FCS

The concept of spot-variation-FCS (svFCS) is to measure diffusion coefficients over a range of spatial scales in order to discriminate between different diffusion modalities [2, 3] (*e.g*., free diffusion, hopping, and, binding). In conventional svFCS, diffusion times are quantified at different detection volume sizes by either (i) changing the pinhole size (in confocal microscopy) or the filling of the objective’s back-aperture [18], increasing the optical complexity, or (ii) by tuning the stimulated emission depletion beam intensity in STED-FCS [19], increasing photodamage. However, in all cases, several sequential measurements are needed, one for each detection volume probed, increasing the experimental time, thus increasing photo-toxicity and hindering the study of fast dynamics. Moreover, sequential measurements limit the study of rapidly evolving systems and are susceptible to artefacts from organelle or cell movements during or in between acquisitions. An alternative strategy to obtain simultaneous measurements is time-resolved STED-FCS, where different effective detection volumes are reconstructed from a single acquisition by sorting photons according to their arrival time after excitation. Progressive time gating reduces the effective observation volume without sequential measurements; however, the signal-to-background ratio decreases markedly for the smallest volumes, limiting the robustness of the correlation analysis [20, 21].

With an array detector, the spot size *ω*_0_ can be chosen in post-processing from a single FFS measurement by summing the signals from different detector elements, Fig. 2 (a).

**Figure 2:**
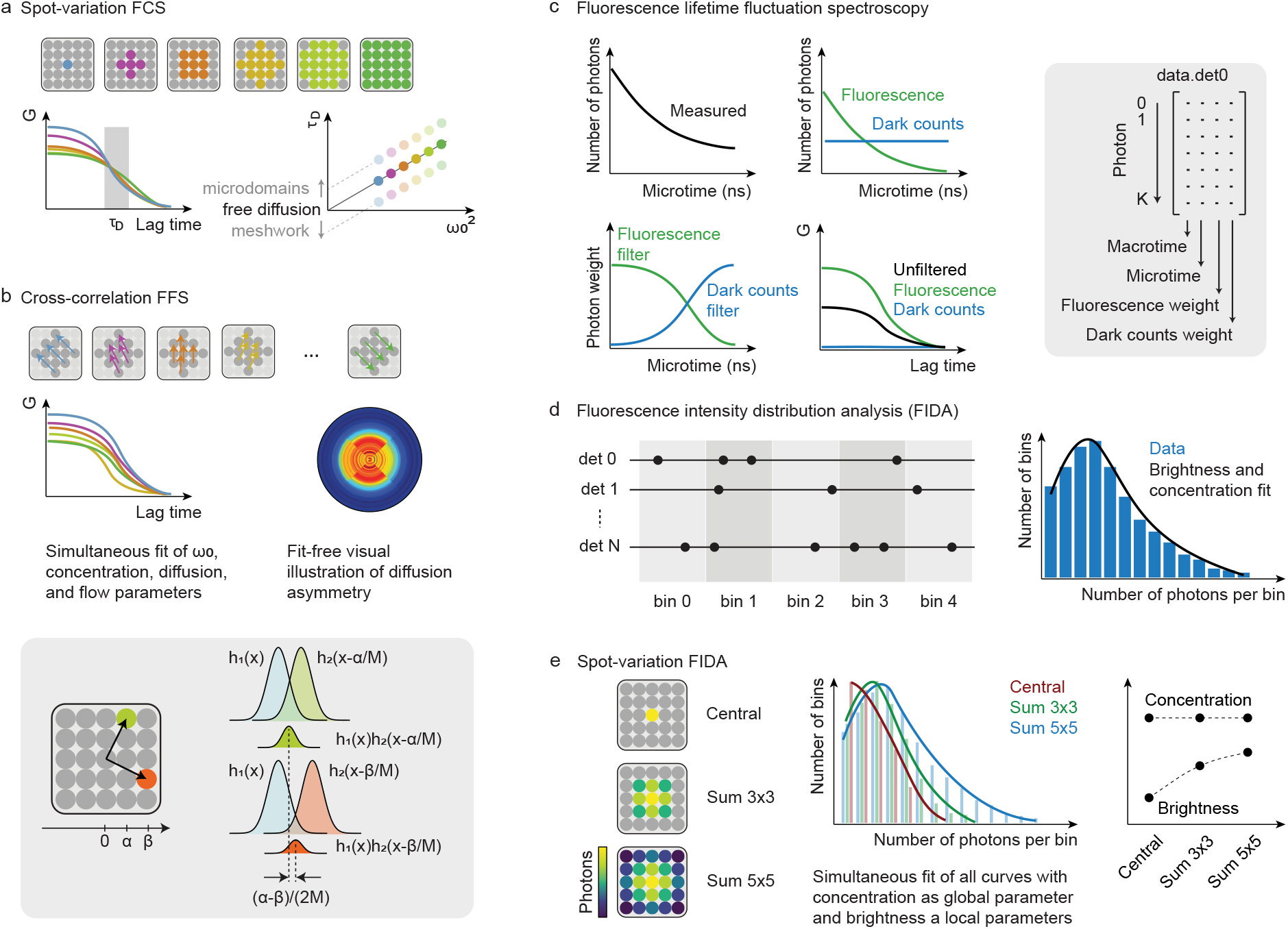
(a) Spot-variation FCS. Summing the signals from different detector elements is equivalent to changing the pinhole size in a confocal microscope, and therefore changing the detection volume. The different correlation curves yield different diffusion times *τ*_*D*_. The relationship *τ*_*D*_ *vs*. the probed area *ω*_2_ indicates the diffusion mode. (b) Cross-correlation FFS. Cross-correlations functions (CCFs) between different pairs of detector elements are calculated and CCFs from pairs with identical shift vectors may be averaged. A global fit yields *ω*_0_, the particle concentration, diffusion coefficient, and flow velocity in the *x,y* direction. In each lateral dimension, the displacement between the focal volumes corresponding to two different detector elements is given by (*α* − *β*)*/*(2*M*), with *α* − *β* the difference in position between the respective detector elements in that dimension and *M* the optical magnification (shaded inset). Plotting four CCFs corresponding to four different directions gives a fit-free visual illustration of potential diffusion asymmetry. (c) Fluorescence lifetime fluctuation spectroscopy to separate different species with a different microtime signature, *e.g*., two components with a different lifetime or a fluorescence and background component. The microtime histogram for a single detector element (top left) is a linear combination of the microtime signature of the two components, here fluorescence and dark counts (top right). By assigning weights to every photon (shaded inset) according to filter functions based on the microtime information (bottom left), filtered correlation curves can be calculated (bottom right). (d) Fluorescence intensity distribution analysis. Photons (black circles) are binned in space and time and the photon counting histogram is fitted to extract the particle brightness and concentration. (e) Spot-variation FIDA. Different histograms can be calculated from differently summed detector elements. If the emitter concentration and brightness are uniform over the detector field-of-view, all histograms can be fitted simultaneously with the concentration as a global fit parameter and the brightness as local fit parameters.

*E.g*., for a 5 × 5 array detector, considering the photons collected by the central element only is equivalent to decreasing the size of the pinhole in a confocal microscope. By considering also the photons (i) in the central 3 × 3 elements or (ii) in all elements, one can increase the virtual pinhole diameter, thus increasing the spot size *ω*_0_. In practice, one can include several more combinations of detector elements, Fig. 2 (a). Fitting the corresponding ACFs yields increasing diffusion times for an increasing number of detector elements considered. For free diffusion, the diffusion time scales proportionally with 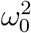 and the diffusion coefficient is inversely proportional to the slope of the curve 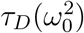. If the diffusion time does not scale proportionally with 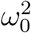, extrapolation of the diffusion law curve 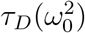 gives information on the type of anomalous diffusion [18]. The *y*-axis intercept 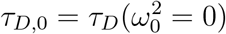 describes the deviation of the diffusion time compared to the expected diffusion time for free diffusion. A positive *τ*_*D*,0_ value indicates domain-confined diffusion while a negative intercept diffusion hindered by a meshwork. Thus, extrapolating 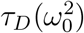 reveals the diffusion modality, a result unattainable with a single conventional FCS measurement. However, isotropically summing the signals from different detector elements causes a loss of information about a potential directionality of the dynamics. To reveal asymmetric diffusion modalities such as flow or diffusion near a border, a more informative method is the analysis of cross-correlations between neighboring detector elements.

### 2.3. Cross-correlation FFS

The different lateral positions of the detector elements can be exploited to detect and quantify directed motion. Consider the case of flow in the positive *x* direction. Then, the cross-correlation curve *G*_*C×R*_ will decay at a later lag time than *G*_*C×L*_, with *G*_*C×R*_ and *G*_*C×L*_ being the cross-correlations between the signals of the central element and its right or left neighbor, respectively, Fig. 2 (b). Assuming that the distance between the focal volumes is known, cross-correlation FFS can reveal concentration, diffusion coefficient, and flow speed in the *x* and *y* directions. Indeed, the cross-correlation function (CCF) for directed motion is:

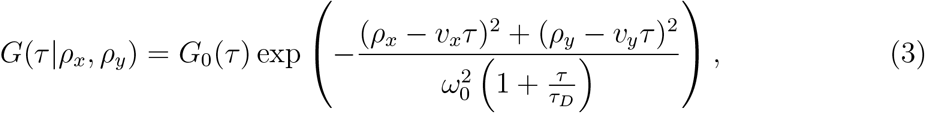

with *G*_0_(*τ*) the ACF from Eq. 2. Here, *ρ*_*x*_, *ρ*_*y*_ are the shifts between the PSFs from the two detector elements. We assume Gaussian PSFs that are identical up to a spatial translation. The paraneters *v*_*x*_ and *v*_*y*_ indicate the flow speed in the *x* and *y* direction, respectively. For an array detector of *N* elements, one can calculate *N* ^2^ CCFs and perform a simultaneous fit of all curves with the diffusion coefficient, *v*_*x*_, and *v*_*y*_ as global fit parameters. Alternatively, to reduce the number of correlation curves and speed up the fit procedure, one may average the CCFs from all detector pairs having the same shift vector ***ρ*** = (*ρ*_*x*_, *ρ*_*y*_). In the case of free diffusion (*v*_*x*_ = *v*_*y*_ = 0), *G*(*τ ρ*_*x*_, *ρ*_*y*_) does not depend on the direction of the shift vector but only on the magnitude 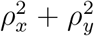. In this case, the number of curves can be reduced further by averaging all CCFs from all detector pairs having the same distance.

However, the assumption of identical Gaussian PSFs for all detector elements is typically not valid. An important advantage of cross-correlation analysis is that it allows for calibration-free measurements, since *ω*_0_ can be kept as a fit parameter, provided that the shift vectors (*ρ*_*x*_, *ρ*_*y*_) for each pair of detector elements are known. The shift vectors can be retrieved from the detector geometry and the total magnification *M* of the optical system. Indeed, consider a Gaussian excitation PSF *h*_1_(**x**) and Gaussian detection PSFs *h*_2_(**x** − **x**_**0**_), with **x**_**0**_ = ***α****/M* and ***α*** the displacement of the detector element with respect to the central element. Assuming no Stokes shift, the total PSF is *h*_1_(**x**)*h*_1_(**x** − ***α****/M*), which is a Gaussian centered at position **x** = ***α****/*(2*M*). Thus, the relative position of the focal volume between any pair of detector elements at positions **x** − ***α*** and **x** − ***β*** is ***ρ*** = (***α*** − ***β***)*/*(2*M*), Fig. 2 (b).

In conventional pair-correlation FCS, spatially separated focal volumes are obtained by moving the excitation beam to different lateral positions [22]. Although this scanning implementation has the advantage that the focal volume is identical for each position in the scanned line, it requires the line time to be shorter than the molecular diffusion time. Instead, with an array detector, multiple positions are probed simultaneously, allowing faster dynamics to be studied.

Direction-dependent diffusion may also result from the presence of barriers, such as the boundary of a large aggregate that prevents molecules inside from diffusing out and *vice versa*. In this case, an FFS measurement near the border will show that the CCFs from detector pairs whose shift vector is perpendicular to the border will be different from CCFs from detector pairs with a shift vector parallel to the border. A model-free, qualitative way to assess direction-dependent diffusion is to plot the CCFs as a polar heat map, where angle encodes direction, radius encodes lag time, and color encodes the CCF amplitude, Fig. 2 (b).

### 2.4. Fluorescence lifetime fluctuation spectroscopy

The combination of FCS and fluorescence lifetime measurement, called fluorescence lifetime correlation spectroscopy (FLCS) [23], is typically realized by combining a single-photon detector with a time-correlated single-photon counting (TCSPC) system. Unlike conventional cross-correlation spectroscopy, FLCS allows cross-correlating signals from two or more types of fluorophores that cannot be spectrally separated. Moreover, FLCS has the advantage that the detection volumes are identical for all species, since only a single excitation beam and detector are used. FLCS can also reveal changes in the fluorescence lifetime caused by changes in the molecular nano-environment, *e.g*., conformational state changes or pH modification.

Fluorescence lifetime fluctuation spectroscopy (FLFS) is the extension of FLCS with an asynchronous (SPAD) array detector. In FLFS, each of the *N*_*c*_ detector elements is connected to a TCSPC data acquisition card, and the raw data consists of *N*_*c*_ lists of photon arrival times. Each arrival time is typically expressed as a combination of a coarse macrotime (*i.e*., the number of laser pulses counted since the start of the measurement) and a fine micro-time (*i.e*., the ps-resolution arrival time of the photon with respect to the last laser pulse). The macrotimes can be used to calculate the ACFs and CCFs while the microtimes give a simultaneous measurement of the lifetime. In addition, similarly to FLCS, correlation analysis of a single component in a mixture and cross-correlation between multiple components is possible by assigning a weight to each photon based on its microtime. Even for a single species, having the arrival time information is useful to correct for detector artifacts such as the dark count rate and the afterpulsing probability. Indeed, the randomness of the dark counts and the long time-scale of the afterpulsing probability lead to an even background in the microtime histogram, Fig. 2 (c). Having a microtime fingerprint that is different from the fluorescence fingerprint means that the detector artifacts can be considered as a second component and one can design a statistical filter to remove this contribution [23, 24, 7]. Artifact filtering can alter the correlation curve in several ways. First, afterpulsing may appear in the unfiltered ACF as an additional fast decay in the microsecond time range and can be misinterpreted as a fluorescence triplet-state contribution. Second, spurious counts lower the ACF amplitude, which leads to an overestimation of the particle concentration from the fitted amplitude. This effect is evident in spot-variation FLFS, where unfiltered ACFs yield a particle-number curve *N* (*V*) that is not proportional to the focal volume *V*, whereas filtering the ACFs shifts the *N* values onto a line passing through the origin [25]. Thus, the extension of FFS with TCSPC is a powerful way to disentangle molecular dynamics from detector-related artifacts.

### 2.5. Fluorescence intensity distribution analysis

The intensity fluctuations measured with FCS can also reveal the apparent molecular brightness, *i.e*., the number of detected photons per molecule per second [26, 27]. For a chosen bin time, a histogram of the photon counts shows a super-Poissonian behavior due to the movement of the molecules through a non-homogeneous illumination and detection profile. In fluorescence intensity distribution analysis (FIDA) [26], the photon count distribution *P* (*n*), *i.e*., the probability of counting *n* photons in a bin, is derived using the representation of generating functions:

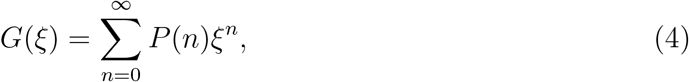

with

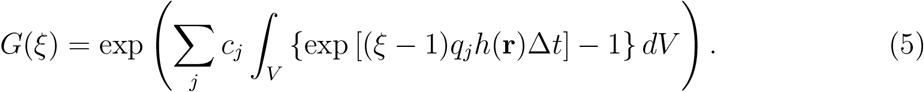

Here, *c*_*j*_ and *q*_*j*_ are respectively the concentration and brightness of component *j, h*(**r**) the PSF, and Δ*t* the bin time. Here, we assume that the deviation from a purely Poissonian distribution is entirely due to particle motion. Other photophysical effects – such as fluorophore blinking, bleaching, and photoselection – as well as detector artifacts – such as dark counts and afterpulsing – can be modeled, but are neglected here.

*P* (*n*) can be calculated by taking the Fourier transform of *G*(*ξ* = exp(*iϕ*)). Since *h*(**r**) is the only term in Eq. 5 that depends on **r**, we can approximate *G*(*ξ*) as follows. First, we discretize *h*(**r**) on a finite 3*D* grid: *h*_*i,j,k*_, with *i, j, k* the index values in the three spatial directions. Let *n*(*A*) denote the number of elements that satisfy condition *A*. Then, we can make a histogram of *h*:

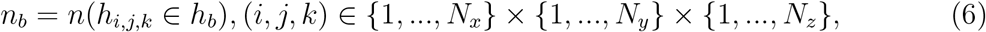

with *n*_*b*_ the number of voxels for which the PSF value lies within the limits of bin *h*_*b*_.

Then,

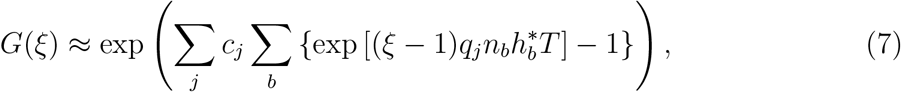

with 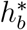 the bin center of bin *h*_*b*_. Eq. 7 can be efficiently computed for any discretized PSF.

With a SPAD array detector, the photon count distribution can be calculated for each element or a combination of elements. *E.g*., in spot-variation FIDA, differently summed combinations of detector elements produce different histograms. If the concentration is not position-dependent, this difference can be entirely attributed to the increased apparent brightness when summing more detector elements. Hence, one can check the reliability of the fit result by checking that all fits produce similar concentration values, and brightness values that scale with the average photon count rate for each detector element. Alternatively, one may fit all histograms simultaneously with the concentration set as a global fit parameter.

## 3. Results

The BrightEyes-FFS software is available as both a Python package on PyPI and GitHub as well as a standalone GUI. For developers or users with at least minimal coding experience, the Python package is recommended as it provides more options and allows custom analysis. The package provides tools for reading FFS data in various file formats, including H5 (BrightEyes-MCS and Genoa Instruments), PTU (PicoQuant), OME-TIFF, and CZI (Zeiss). For intensity data stored in the H5 format, the software expects a 2D array [*N*_*t*_ × *N*_*c*_] stored with the key *data, N*_*t*_ the number of time points, and *N*_*c*_ the number of detector elements. Data collected with BrightEyes-MCS [28] is by default stored in this format. For TCSPC data stored in the H5 format, the software searches for attributes called *detx*, with *x* the detector element number, containing a 2D array with the macro- and microtimes of the photon arrivals in picoseconds. Fig. 3 shows an examplary data set analysed with the Python package.

**Figure 3:**
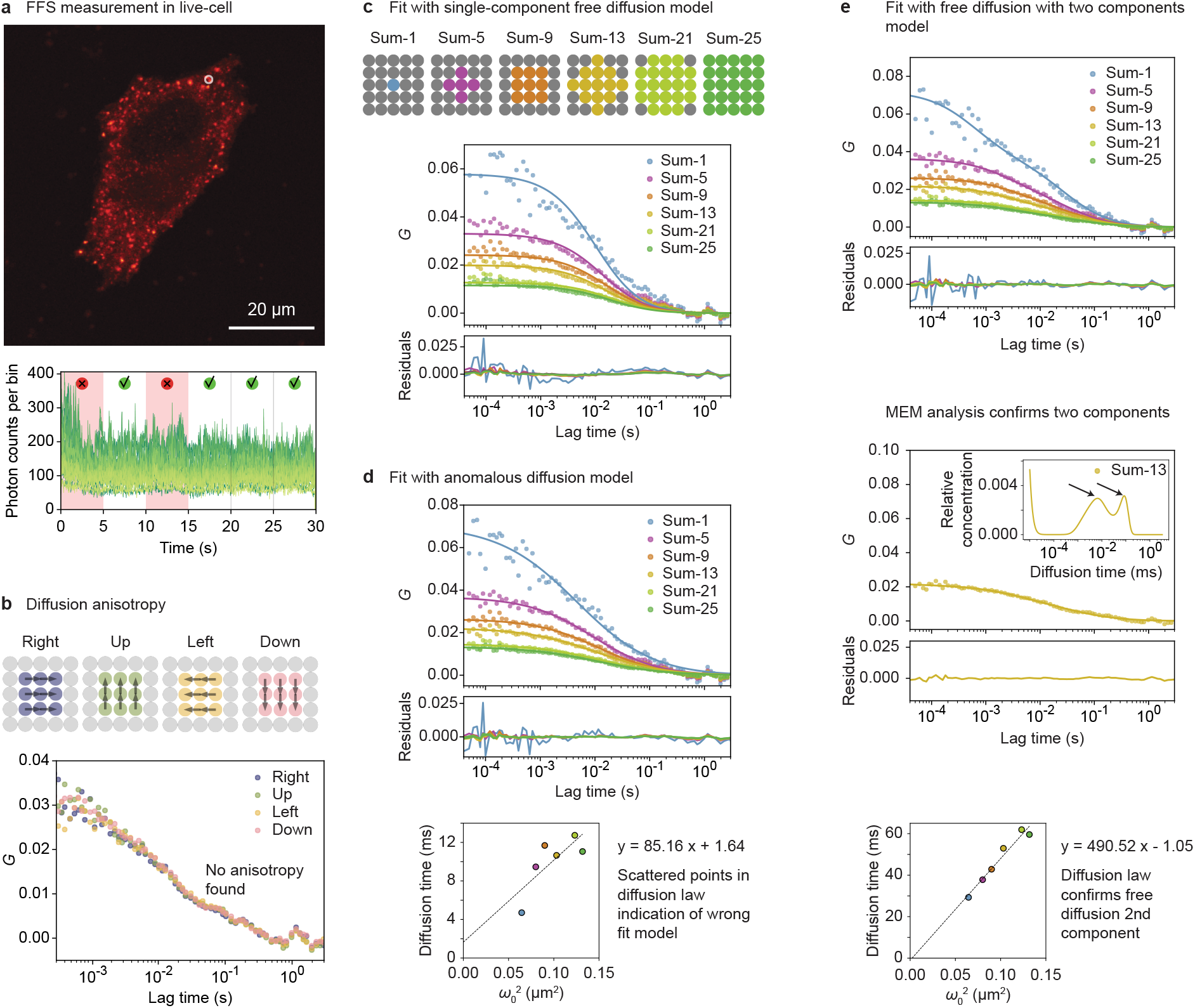
Example of FFS analysis in a living SK-N-BE cell WT expressing G3BP1-eGFP during oxidative stress. Stress granules are visible in the cytoplasm. Details on the sample in [8]. (a) Confocal image indicating the position of the FFS meausurement (top) and fluorescence intensity time trace for each of the 25 detector elements (bottom). Correlation were calculated in segments of 5 s. Two segments (red shades and icons) were detected as outliers and excluded from further analysis. (b) Cross-correlation analysis shows no direction-dependent diffusion. (c) Spot-variation FFS curves and fit with a 1-component free diffusion model. (d) Spot-variation FFS curves and fit with a 1-component anomalous diffusion model. The scattered points in the diffusion law suggest that the curves were fit with the wrong model. (e) Spot-variation FFS curves and fit with a 2-component free diffusion model (top). Example of MEM analysis on one of the correlation curves, showing two peaks in the diffusion time histogram (arrows), suggesting the presence of two components (middle). The corresponding diffusion law for the slow component shows free diffusion.

Experimental artifacts, such as fluorescent aggregates or the erratic movement of living cells, can significantly degrade correlation quality. Consequently, a standard preprocessing strategy involves partitioning the intensity signal into discrete segments for individual analysis, allowing for the targeted removal of those compromised by artifacts. Following this concept, the BrightEyes-FFS software allows segmenting the data into chunks, with ACFs and CCFs calculated for each segment. Correlations can be calculated in either the time domain, the frequency domain using the Wiener-Khinchin theorem, or in time-tagging mode [16]. Segments with low-quality correlations can be automatically detected [29] and excluded from further analysis, Fig. 3 (a). The ACF for any custom combination of detector elements and the CCF between any two detector elements or combinations thereof, can be calculated, allowing the software to adapt easily to custom detector configurations or analysis modalities. Common analysis modalities, such as sv-FFS and cross-correlation FFS, can be accessed through predefined keywords. In this way, the code provides flexibility for custom analyses while remaining user-friendly for routine use.

BrightEyes-FFS includes a variety of analytical models for fitting correlation curves, including models that account for SPAD-specific artifacts at short lag times, such as afterpulsing. The software contains several options to study mobility from free diffusion to directionally asymmetric diffusion, including calibration-free pair-correlation analysis and cross-correlation for flow analysis, Fig. 3 (b-e). Multiple curves can be fitted simultaneously, with the option to treat each parameter as either a global (i.e., identical across all curves) or individual parameter. Fitting is performed by minimizing a weighted sum of the squared residuals using the *least squares* function from the SciPy.optimize library, with the weights *w*(*τ*) equal to the inverse of the variance of *G*(*τ*), i.e., *w*(*τ*) = 1*/σ*^2^(*G*(*τ*)), calculated over the different segments. In addition, we implemented the diffusion asymmetry heat map as a quick and qualitative check for flow or barriers.

For TCSPC data, the package allows filtering the correlation curves based on the microtimes [23, 25]. This method gives a weight to every photon based on its arrival time and allows separating the correlation functions from different components with a different arrival time signature, such as two fluorescent species with a different lifetime, or removing parasitic contributions such as afterpulsing.

Fluorophore concentration and brightness can be retrieved via fluorescence intensity distribution analysis (FIDA). Similar to FFS, histograms for any combination of detector elements can be calculated. We implemented the numerical evaluation of the integral of Eq. 5 to fit the histograms, and therefore no analytical approximation of the PSF shape is needed. The user can input either a 3D array for the PSF, *e.g*., from a z-stack measurement of fixed fluorescent beads or a simulation [17], or a set of two numbers [*ω*_0_, *z*_0_*/ω*_0_], with *ω*_0_ and *z*_0_ the width and height of a PSF approximated as a 3*D* Gaussian function. Fit functions for any number of species are available, and SPAD detector artifacts such as the dark count rate are included in the model. The user can choose among different loss functions, such as the sum of squared errors or the negative log-likelihood.

For scientists with minimal software development experience, no or minimal coding is required. The BrightEyes-FFS GUI can be downloaded as a Windows executable from Zenodo. The GUI is built on the Python BrightEyes-FFS package and is available as a standalone file than can run without installation. A schematic of the GUI is presented in Fig. 4. The user can load an image of the sample (*e.g*., a cell) and multiple single-point FFS measurements (*e.g*., taken at different positions in the same cell) for analysis. Most features of the BrighEyes-FFS package are available in the GUI. The GUI software is also open-source and can, with minimal coding requirements, be adapted for custom analysis (*e.g*., for adding custom fit models). In addition, the GUI has one-way integration with Jupyter Notebook; a session started in the GUI can be continued in Jupyter Notebook for custom plotting or further analysis. To support users with limited programming experience, the GUI can automatically generate notebooks. These notebooks allow users to view analysis results or replicate the entire analysis workflow from scratch by simply executing the provided code, eliminating the need to write custom scripts. Moreover, several exemplary notebooks are available on the GitHub page of the project, as well as a Wiki with guidelines and documentation on the most important features of the library.

**Figure 4:**
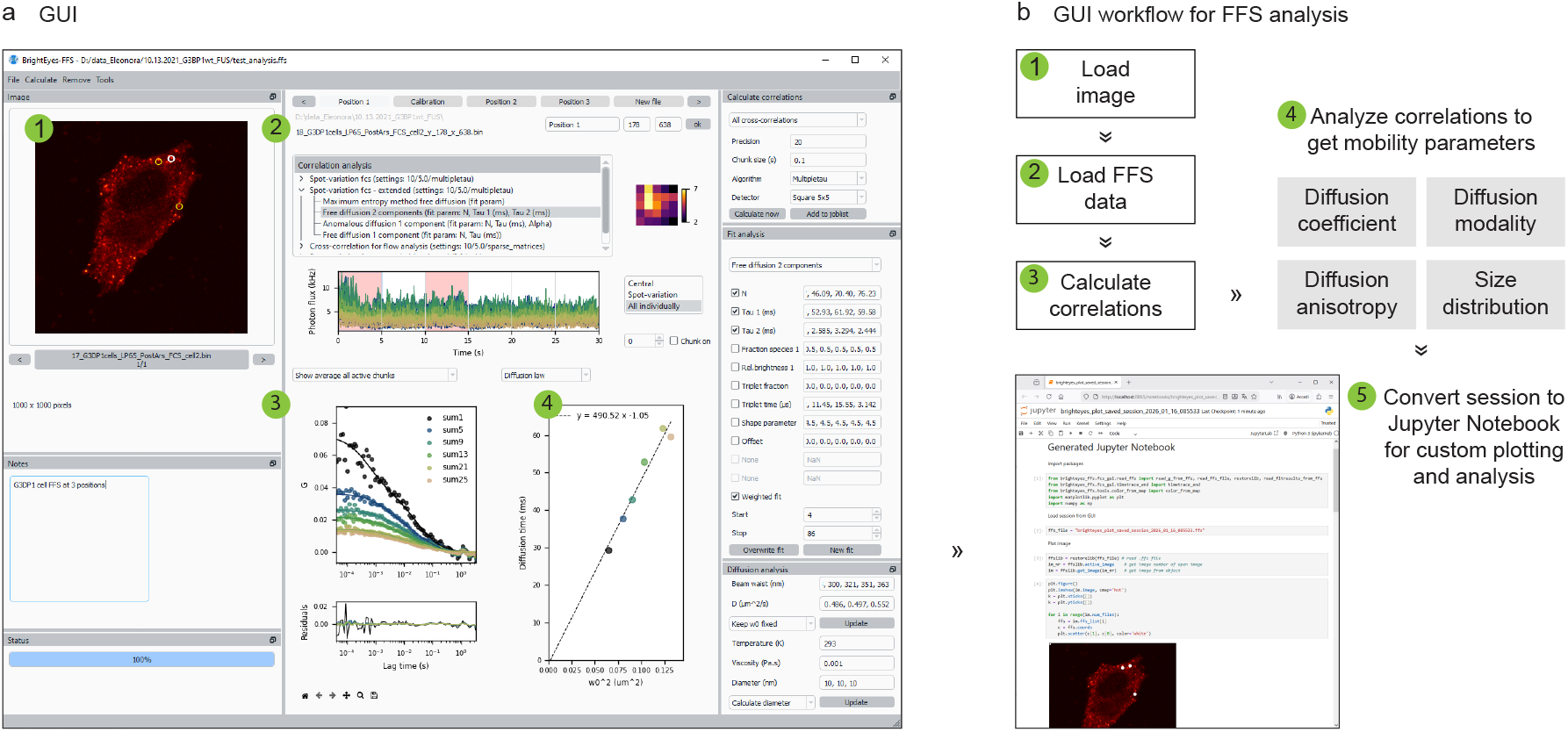
(a) Graphical user interface (GUI) for FFS analysis, showing a cell image and different locations where FFS measurements were done. (b) Recommended workflow for FFS analysis.

## 4. Conclusions

In summary, BrightEyes-FFS is an open-source Python-based environment for FFS analyis. While currently optimized for square SPAD array detectors (*e.g*., 5 × 5 or 7 × 7 pixels) such as the PRISM-Light detector by Genoa Instruments, the software can also open *PTU* files from the PicoQuant Luminosa PDA-23 array detector and *CZI* files from the Zeiss Airyscan detector. Indeed, regardless of the spatial arrangement of the detector elements, any FFS data set can be interpreted as either a 2D array [*N*_*t*_ × *N*_*c*_] with *N*_*t*_ the number of time points and *N*_*c*_ the number of detector elements or a 2D array [*N*_*p*_ × 3] containing for each of the *N*_*p*_ photons the macrotime, microtime, and detector element number. Modifications such as summing over different detector element numbers or calculating cross-correlations between different detector elements, as well as adding new fit models, can be done with minimal coding. In addition, we anticipate integrating BrightEyes-FFS as an add-on module to the microscope control software, providing realtime data feedback to enhance user efficiency in data collection. An important step in this direction is the Python interface recently released by PicoQuant for Luminosa, called

LumiPy, which allows the BrightEyes-FFS to be imported in the microscope control software. Thus, we hope BrightEyes-FFS will help users from various scientific backgrounds to do FFS, thereby helping to unravel the complex dynamics of life.

## Data availability

No new data are presented in this manuscript. The data shown in Fig. 3 was taken from [8].

## Code availability

The source code of BrightEyes-FFS is available on GitHub (https://github.com/VicidominiLab/BrightEyes-FFS) and on the Python Package Index (https://pypi.org/project/brighteyes-ffs). The source code of the BrightEyes-FFS GUI is available on GitHub (https://github.com/VicidominiLab/BrightEyes-FFS-GUI). A standalone executable of the GUI, with an exemplary data set, is available on Zenodo (https://doi.org/10.5281/zenodo.14826849). Queries and feedback can be provided by creating an issue on GitHub.

## Acknowledgements

The authors thank Stijn Dilissen and Jelle Hendrix (Hasselt University, Belgium) for providing FFS data with the Zeiss Airyscan, and PicoQuant for providing FFS data with the Luminosa system. The research was supported by: the European Research Council, BrightEyes No. 818699 (E.S., G.V.) and MotionPicture No. 101219234 (ES); the National Recovery and Resilience Plan (NRRP), Mission 4, Component 2, Investment 1.1, Call for tender No. 1409 published on 14.9.2022 by the Italian Ministry of University and Research (MUR), funded by the European Union - NextGenerationEU - Project Title:ALPHA (202288M84C) - CUP D53D23001060006 (G.V.).

